# A quantitative model for metabolic intervention using gut microbes

**DOI:** 10.1101/2020.04.01.020677

**Authors:** Zachary JS Mays, Nikhil U Nair

**Affiliations:** Department of Chemical & Biological Engineering, Tufts University, Medford, MA 02155

## Abstract

As medicine shifts toward precision-based and personalized therapeutics, utilizing more complex biomolecules to treat increasingly difficult and rare conditions, microorganisms provide an avenue for realizing the production and processing necessary for novel drug pipelines. More so, probiotic microbes can be co-opted to deliver therapeutics by oral administration as living drugs, able to survive and safely transit the digestive tract. As living therapeutics are in their nascency, traditional pharmacokinetic-pharmacodynamic (PK-PD) models for evaluating drug candidates are not appropriate for this novel platform. Using a living therapeutic in late-stage clinical development for phenylketonuria (PKU) as a case study, we adapt traditional oral drug delivery models to properly evaluate and inform the engineering of living therapeutics. We develop the <u>a</u>dapted for <u>l</u>iving <u>t</u>herapeutics <u>c</u>ompartmental <u>a</u>bsorption and <u>t</u>ransit (ALT-CAT) model to provide metrics for drug efficacy across nine age groups of PKU patients and evaluate model parameters that are influenced by patient physiology, microbe selection and therapeutic production, and dosing formulations.

**Importance:** This work describes a kinetic model to study the behavior of orally delivered living therapeutics. Such therapeutics are becoming increasingly relevant and are an exciting mode of drug delivery that stems from the growing interest through the convergence of advances in synthetic biology of probiotics and gut microbes as well as microbiome science. In particular, this work describes the development of a mathematical framework (pharmacokinetic-pharmacodynamic, PK-PD) called ALT-CAT to model the behavior of orally delivered engineered bacteria that act as living therapeutics by adapting similar methods that have been developed and widely-used for small molecular drug delivery and absorption.

## Introduction

The most common and practical route of drug delivery is oral ingestion, pairing patient convenience and high compliance with ease of administration and stability. While a simple initial assessment of a drug’s biochemical feasibility for oral delivery can be predicted using the biopharmaceutical classification system (BCS) (1) or rule of five (Ro5) (2), these evaluations do not inform the further complexities of selecting lead candidates, developing a dosage or release strategy, effective bioavailability, and pharmacokinetics. Accordingly, a range of models – *in vitro* (3) and *in vivo* (4), static (5) and dynamic (6, 7), molecular (8) and system environment (9) – are used extensively in drug development to evaluate and hone desired features.

An economically appealing alternative, trading human capital and consumables for computational resources (time and power), is the modeling of drug absorption in the gastrointestinal tract (GIT). *In silico* models have been developed for both molecular interactions (10), such as receptor binding or transport, and physiology-based pharmacokinetics (PBPK) (11), such as compartment disposition or clearance. Several commercial software packages based on these models (12–14) incorporate drug physiochemical properties (solubility, degradation, permeability, molecular size, aggregation, charge), formulation properties (dosage, drug release profiles, absorption enhancers, matrix polymorphism), and GI physiological properties (gastric emptying, intestinal pH, motility, luminal content, transporters, metabolism, epithelial sequestration, disease state). Most notably, GastroPlus™ (Simulations Plus, Lancaster, CA) was conceived from the advanced compartmental absorption and transit (ACAT) model (15), separating the small intestine (SI) into a series compartments and solving the differential equations surrounding drug mass transfer.

While *in silico* oral drug absorption models have proven invaluable to the drug development pipeline, an emerging class of therapeutic candidates are not addressed with current methodologies. Living therapeutics (also termed microbiome-based therapeutics, microbial biotherapeutics, and probiotics 2.0) are engineered microorganisms that have been programmed to produce drugs, potentially with autonomous contextual awareness (16–18). These microbes are chosen and designed to be compatible with the human gut microbiota, acting not only as drug producers but as delivery vehicles that can survive and traverse the harsh environment of the gut. The living therapeutics platform offers several potential advantages including a constant or timed drug release, real-time dose changes, spatial targeting of delivery, sub-clinical detectability, and the production of large or complex biomacromolecules normally incompatible with oral delivery (19). The latter is the basis of an enzyme-substitution therapy (EST) for phenylketonuria (PKU) in probiotic *Escherichia coli* Nissle 1917 (EcN) that is currently in Phase 1/2a clinical trials (SYNB1618, Synlogic Inc; ClinicalTrials.gov identifier NCT03516487) (20). Here, the enzyme phenylalanine ammonia-lyase (PAL) expressed in EcN converts excess toxic phenylalanine (phe) to the non-metabolite *trans*-cinnamic acid (*t*CA) and provides therapeutic benefit.

It is clear where the limitations of current oral drug absorption models arise when considering living therapeutics. The “drug” being delivered is actually a microbe that produces a catalytically active enzyme, can adhere to the gut mucosal layer, can replicate, and change PAL expression levels. Furthermore, if a PKU patient consumes protein containing phe, absorption of phe is counterproductive, while degradation (i.e. PAL conversion of phe to *t*CA) is the desired outcome. Thus, this model must account for competition between absorption of a toxin (phe) and catalytic depletion by an enzyme. Finally, just as first-pass hepatic elimination affects a drug bioavailability profile, enterorecirculation of amino acids can affect concentrations of phe in the gut lumen and boost the efficacy of PAL independent of meal phe (21).

Using a PAL-based treatment of PKU as the case study for living therapeutic treatments (20, 22–25), this work presents the <u>a</u>dapted for <u>l</u>iving <u>t</u>herapeutics compartmental <u>a</u>bsorption and <u>t</u>ransit (ALT-CAT) *in silico* model, which incorporates the physiology of the GI tract, different bacterial chassis, and different expression levels of PAL into the traditional compartmental model (9). The CAT model was adapted to populate the SI with a microbial population and combined with the stochastic gastric emptying (sGE) model to simulate a non-linear emptying of the stomach. A fraction of the microbial population was designated to be a living therapeutic expressing PAL to degrade phe. A therapeutic factor (TF) was created to assess model parameters such as bacteria population, enzyme activity, and phe transport. Target concentrations of phe in the blood were used as output for therapeutic efficacy. This work presents a refreshed toolset for the evaluation and comparison of living therapeutic constructs and their parts therein, serving as a benchmark for designing orally administered drug formulations.

## Model construction

The original CAT model determined that a cumulative time frequency distribution of 400 human SI transit times is best represented by seven equal transit compartments (Equation 1). Drug absorption can then be derived within each compartment, relating drug bioavailability to its effective permeability coefficient (*P_eff_*) (Equation 2). Drug dissolution, pH-dependent solubility, first-pass hepatic metabolism, degradation, and other region-dependent factors were added later in the ACAT model, the foundation for GastroPlus™ software. With these equations, *M_n_* is the amount of drug in the n^th^ compartment (mg); *k_T_* is the transit rate constant through the compartment (min^-1^); *T_SI_* is the average transit time through the small intestine (min); *r* is the luminal radius (cm); and *F_a_* is the fraction of dose absorbed (9). In our modification, we model the dynamics of toxin (phe in case of PKU) absorption using these equations.

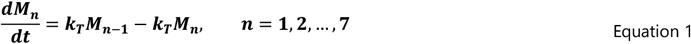

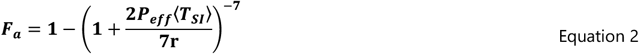

While gastric and colonic drug absorption were included in the CAT model as well, the stomach is modeled by linear emptying, and flow is estimated by first-order ordinary differential equations as a plug-flow reactor (26). Physiologically, gastric emptying is a more complex process involving churning contractions, both peristalsis and retroperistalsis, controlled opening of the pyloric sphincter, solid versus liquid digesta, and hormonal feedback to macronutrient composition. These coordinated movements are observed as sporadic and irregular bolus injections over approximately three to four hours. Several continuous smooth equations have been employed to loosely generalize observed data (27), but more recently, to better account for this, a stochastic gastric emptying (sGE) model was developed by adapting the power exponential form of a standard Wiener process (Equation 3). This was then used to track a decreasing monotonic jump process (Equation 4). Here, *α* and *β* are empirical parameters found to give 90% confidence intervals that describe the data in Locatelli, *et al.* (2009) (28).

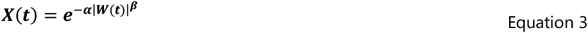

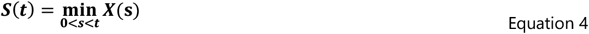

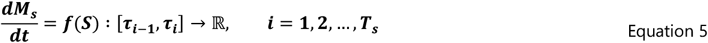

A crucial difference between modeling traditional oral drug delivery and an EST is that the former is designed to maximize absorption (and therefore bioavailability) of a drug, while the latter is meant to prevent absorption of a toxic metabolite. With highly permeable or quickly transported small molecules, such as phe in the case of PKU, the rate limiting step of absorption is gastric emptying. As such, modeling gastric emptying more accurately with the sGE model and combining this with the ACAT model was the framework for constructing the ALT-CAT model for PKU treatment (Figure 1). A spline function can be interpolated from the sGE model over the total gastric emptying time (*T_GE_*) to calculate instantaneous emptying rates entering the SI (Equation 5).

**Figure 1.**
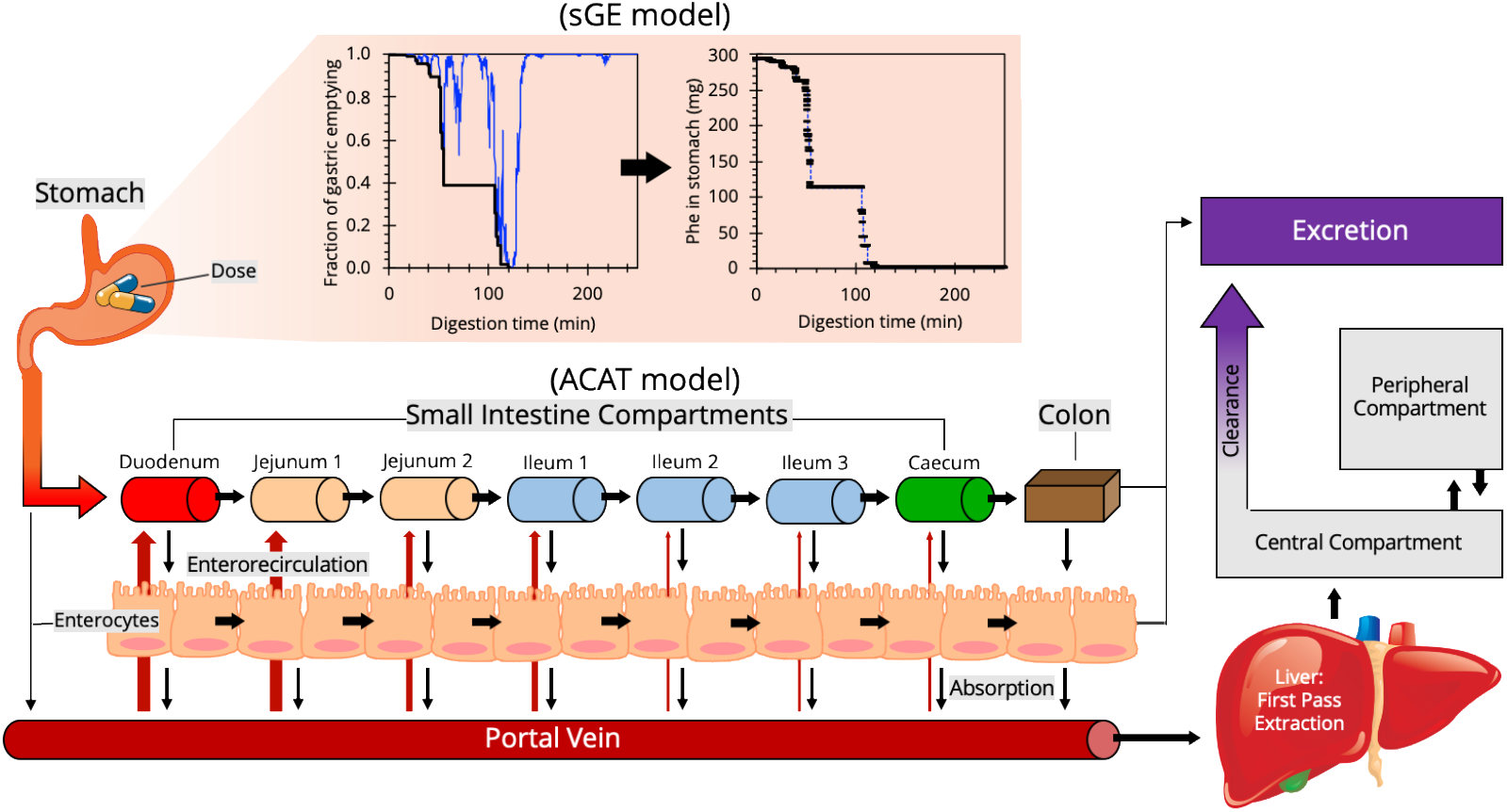
Schematic of the ALT-CAT model for drug absorption. As in the ACAT model, the stomach empties into a seven compartment SI. Mass transfer differential equations surround the fate of a drug in each compartment, which ultimately defines drug bioavailability. The ALT-CAT model adapts this framework by incorporating the stochastic gastric emptying (sGE) model (orange) of the stomach and, in the case of PKU, models phe absorption as detrimental and phe degradation as therapeutic, with enterorecirculation (red arrows) contributing to luminal phe. Adapted from Agoram et al. (2001) and Macheras et al. (2003) (15, 29).

Within each compartment of the ACAT model, the instantaneous concentration of a drug transiting the gut (30) is balanced with diffusion or transport into the plasma, dissolution, degradation, aggregation, and pH-dependent solubility. While certain physiochemical or formulation properties are simplified with phe, the ALT-CAT model requires the additional mass transport of phe from the blood back into the gut lumen, a phenomenon known as amino acid enterorecirculation or exsorption. This likely occurs from both epithelial polarity (21) – active and passive transport variation between the apical (*AP*) and basolateral (*BL*) membranes of the endothelium – and pancreatic and glandular secretions during digestion (30). Transport of phe across a Caco-2 monolayer has been previously modeled as a combination of saturable and non-saturable uptake (*u*) and efflux (*e*) in both directions (Equation 6-8) (31), approximately ten times higher in the AP-to-BL direction (32). Nonlinear kinetics require phe-specific max flux (*J_max_*) (mmol mg^-1^ min^-1^) and half maximum concentration (*K_M_*) (mM) values, while first-order kinetics are described by phe-specific flux constants (*f*). Predicted plasma (*p*), intracellular (*i*), and luminal (*l*) phe concentrations can then inform the model, and enterorecirculation is represented as a net flux (*J_net_*) ratio and incorporated in the n^th^ compartment absorption scale factor (*β_n_*).

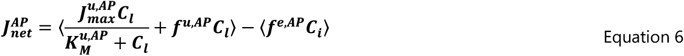

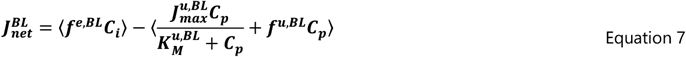

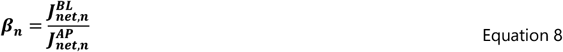

To complete the ALT-CAT model, phe degradation can be reframed as therapeutic conversion by PAL. The GIT is populated with more than 1000 species of microorganisms in a community known as the human microbiota. This diverse population is thought to have evolved with the mammalian gut with spatial organization and niche regioselectivity determined by inextricably complex community interactions and ecological synergy. While much of the human microbiota reside in the large intestine, a low but longitudinally increasing microbial population can survive and reside in the SI. The boundaries for microbial biomass present in the duodenum, jejunum, and ileum have been reported (33), with lactic acid bacteria, a group in which the majority of probiotics are classified, encompassing approximately 0.1-10% of the population (34). Using these regional population metrics, the SI can be randomly populated with a living therapeutic population (*M_LT_,n)* that defines n^th^ compartment PAL delivery (Figure 2). Furthermore, phe conversion (i.e. PAL expression and activity) can be approximated across the population, with Michaelis-Menten enzyme kinetics (*J_max_* and *K_M_)* defining both phe transport (Equation 9) and PAL activity (Equation 10). These adjustments form the ALT-CAT model (Equation 11), using changes in blood plasma phe concentration (Equation 12) as a metric for therapeutic efficacy.

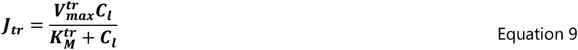

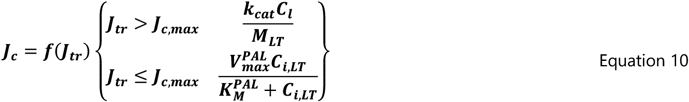

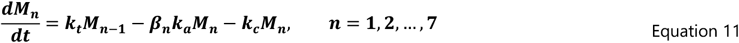

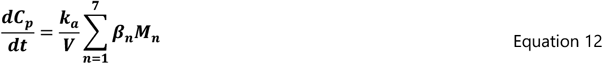

**Figure 2.**
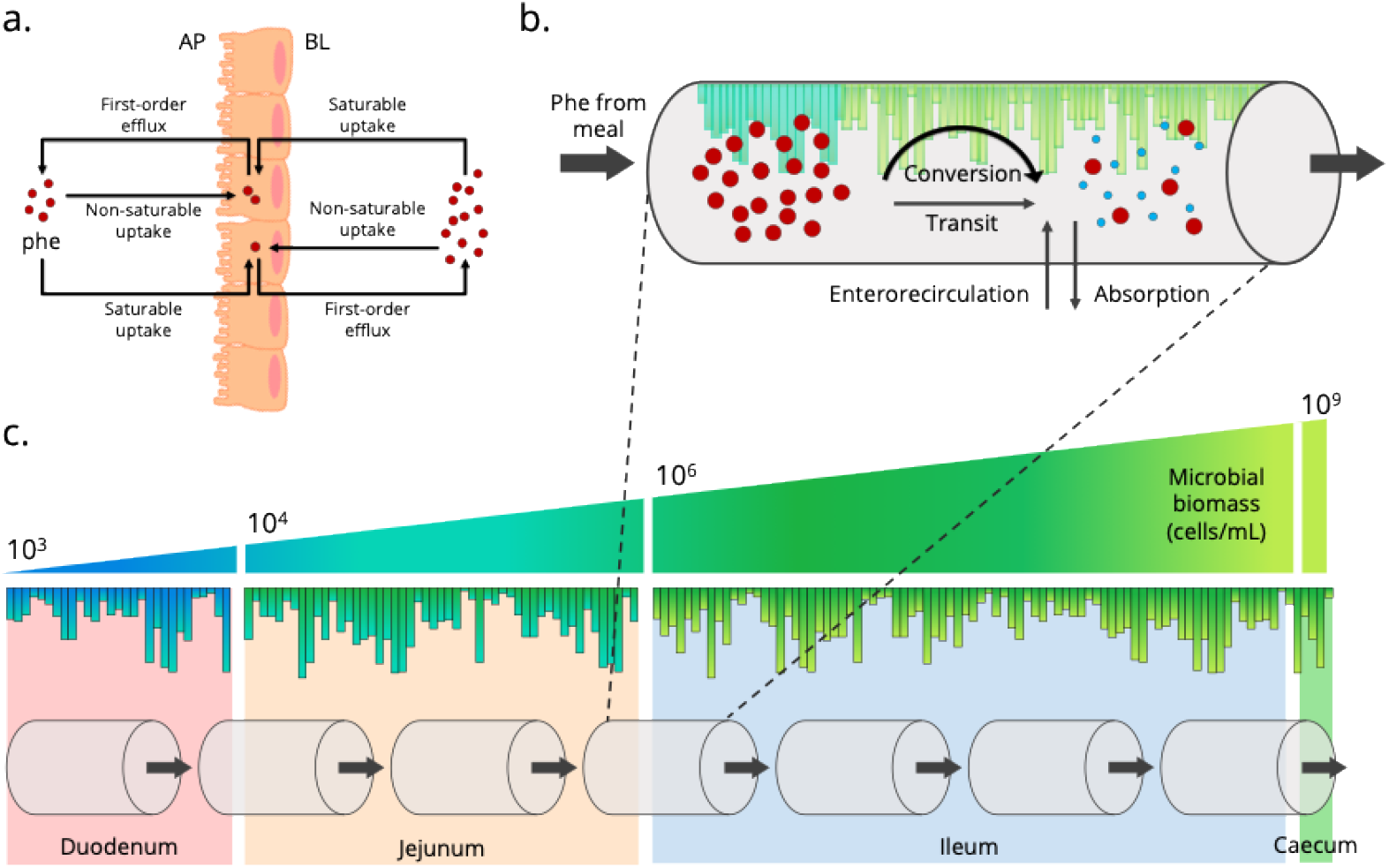
Populating the small intestines with living therapeutics (a.) In PKU, enterorecirculation of phe back into the lumen contributes to (b.) the mass transfer equations driving phe degradation (therapeutic conversion) by PAL present in each compartment’s living therapeutic population. (c.) Each section of the SI has a defined microbial population range. Cross-sections, similar to column plate counts in chromatography, can be generated based on gut length, each with a bounded random microbial population. The full SI can then be equally sectioned into seven compartments as in the ACAT model. Enterorecirculation adapted from Hu and Borchardt (1992) (30). The MATLAB files for model can be found online (https://github.com/nair-lab/ALT-CAT)

## Results & Discussion

Solutions to the differential equations of the ALT-CAT model can be visualized as the change in the phe in each compartment as compared to the start-of-day plasma phe. An example average adult PKU patient, eating four meals of low phe protein over the course of the day, exhibiting 0.8% living therapeutic coverage, is represented in Figure 3. As each meal empties into the SI, it creates a compartmental spike in phe the reverberates down the SI. Blood plasma phe slowly increases over the day, peaking at midnight with a 195% increase and does not follow gastric emptying, indicating the impact that enterorecirculation has on therapeutic conversion. This is similarly highlighted by the high concentration of degraded phe (i.e *t*CA) in the urine.

**Figure 3.**
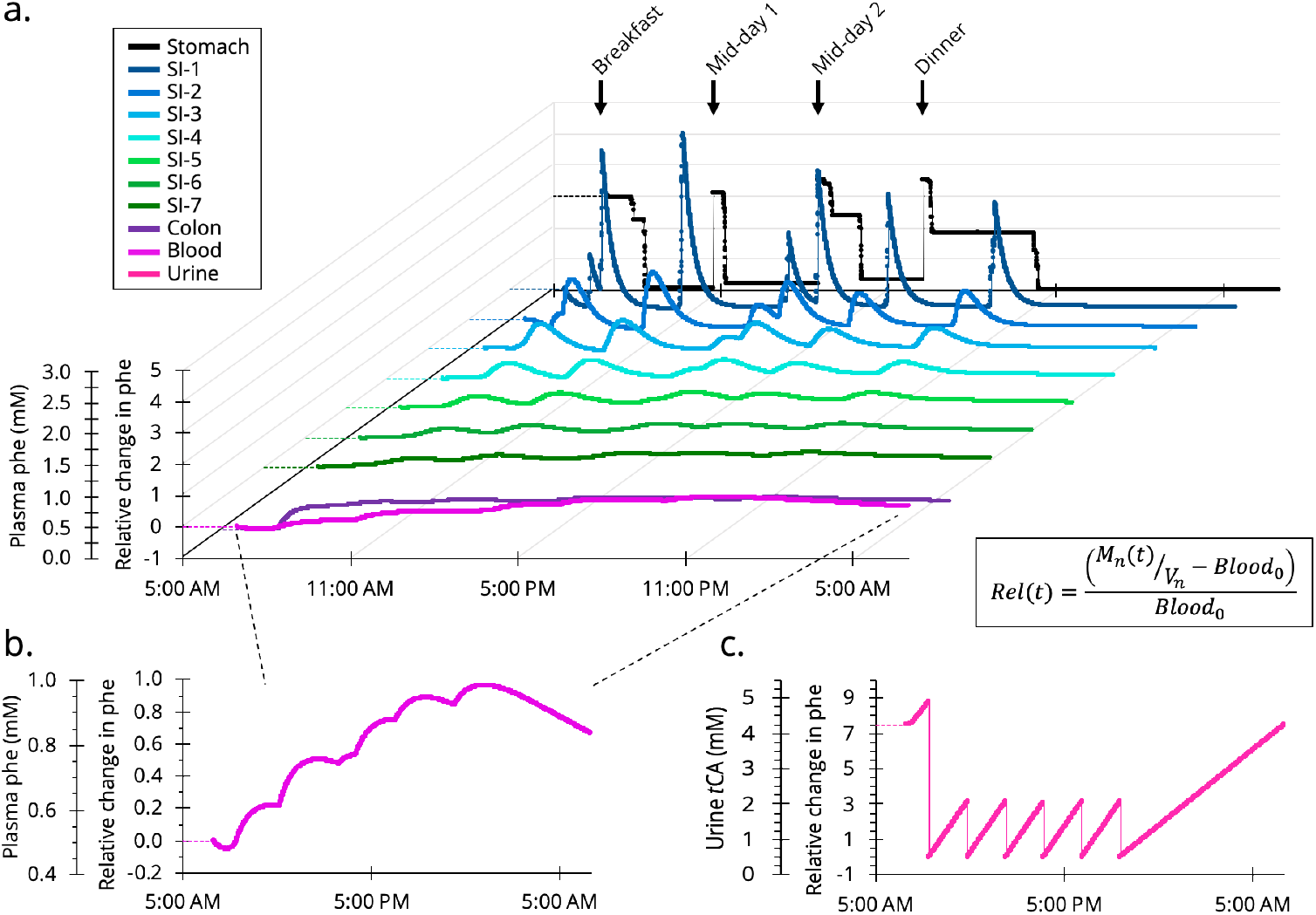
Representative solution to phe digestion in an adult using the ALT-CAT model. (a.) The sGE model generates a release schedule of phe for 4 meals (average adult) over the course of 24 h. Differential equations quantify instantaneous phe in each compartment, visualized as the relative change (percent difference) in phe compared to a start-of-day plasma phe of 0.5 mM. (b.) Blood plasma phe varies by phe absorption and exsorption while a population of living therapeutic converts phe in the lumen of the seven SI compartments. (c.) Urine excretion of *t*CA, or an equivalent biomarker such as hippurate, is proportional to PAL specific activity.

Several physiological parameters of the ALT-CAT model can drastically change across a population, and thus inform, the efficacy of a living therapeutic (Table 1). Drug absorption is proportional to the *P_eff_*, which typically defines the bioavailability of large complex drugs with poor solubility. In the case of PKU, phe is an essential amino acid that is readily absorbed without much resistance, observed as sequentially less phe present in compartments of the ALT-CAT model (Figure 3). Physiological properties that change between age and sex groups, such as the SI length and diameter, dictate the available surface area for absorption, with villi and microvilli amplifying the cylindrical surface area by 60–120 times (35). Additionally, when considering a metabolic disorder such as PKU, body size, amount of protein per meal, number of meals per day, frequency of feeding, and sleep schedules can all drastically impact the availability, and thus metabolism and therapeutic degradation of phe.

**Table 1.**
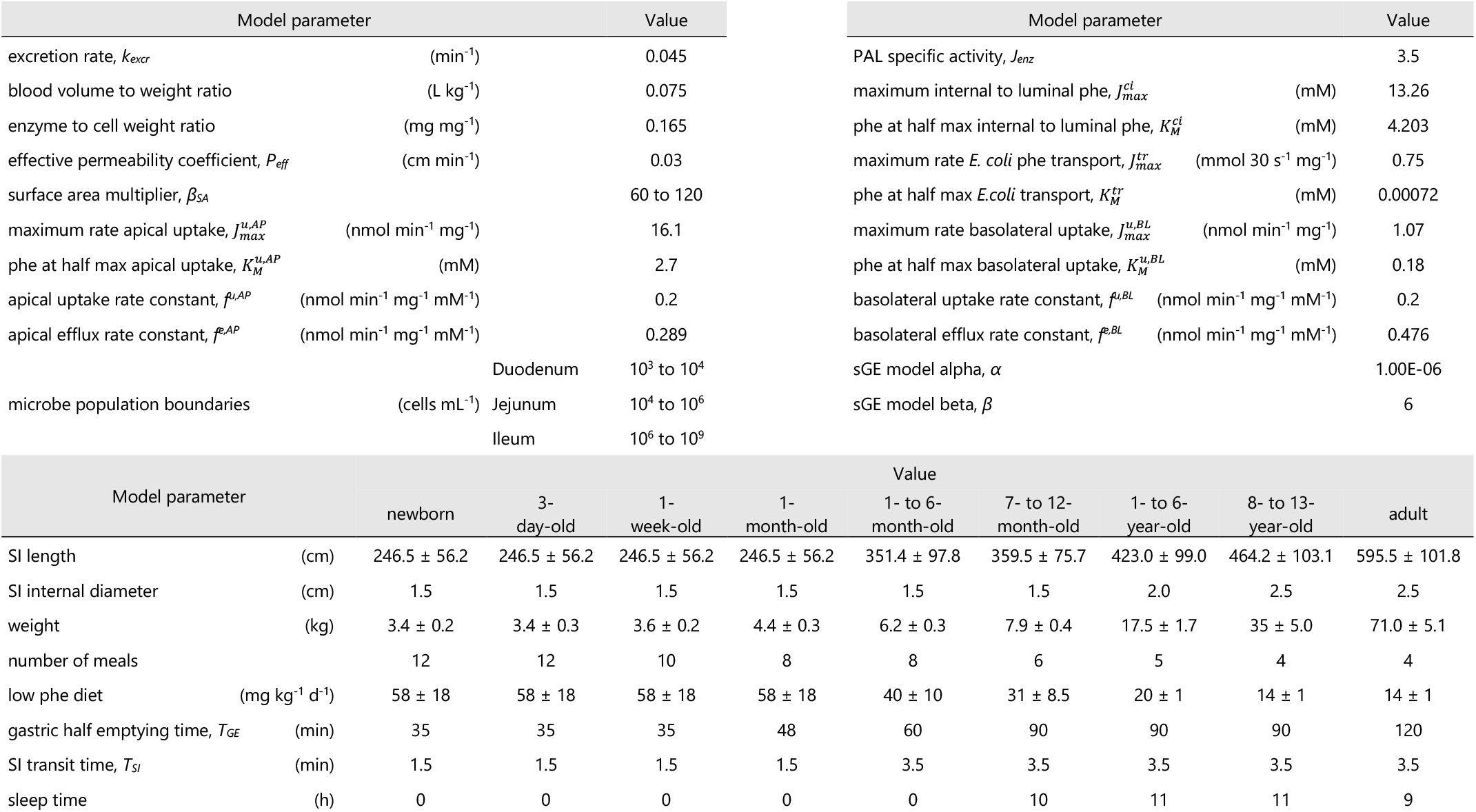
Parameters entered into the ALT-CAT model

We ran nine age groups through the ALT-CAT model, chosen based on physiological properties of the stomach and SI. Starting with a medium-high blood plasma phe concentration (0.5 mM) and 1% living therapeutic population, the end-of-day concentration for 1000 simulations were observed to be influenced by these physiological factors (Figure 4). Each simulation is randomly built using a normal distribution of gut length, body weight, meal phe content, surface area multiplier, and microbial population (Table 1). For the infant and toddler groups, there was much more variability in outcomes. Notably, an all-liquid diet at this age is better modelled as a linear gastric emptying process, which, alongside a large number of meals and short intestine lengths, leave little time for therapeutic conversion of phe. With the majority of phe being freely absorbed, the likely significant contributor to the final distributions is the surface area multiplier (*β_SA_*). Perhaps an adjuvant strategy involving phe transport inhibitors would be impactful for these age groups. Large neutral amino acid (LNAA) supplementation has been effective in PKU adults (36), though with the importance of amino acid concentrations for neural development, not currently recommended for children. Inhibition of epithelial LNAA transporters, such as T-type amino acid transporter (TAT1), with D-isomers or phe derivatives could more specifically target intestinal absorption (37, 38). The ALT-CAT model reiterates reported evidence that gastric emptying is rate-limiting, and once the sGE model is implemented with 7- to 12-month-olds, slower phe introduction corresponds to better therapeutic conversion (i.e. lower plasma phe concentration). Older age groups have designated hours of long sleep as well, allowing phe to be degraded without a meal dictating concentrations. Larger GITs obviously provide more space for more microbes as a percent of the population.

**Figure 4.**
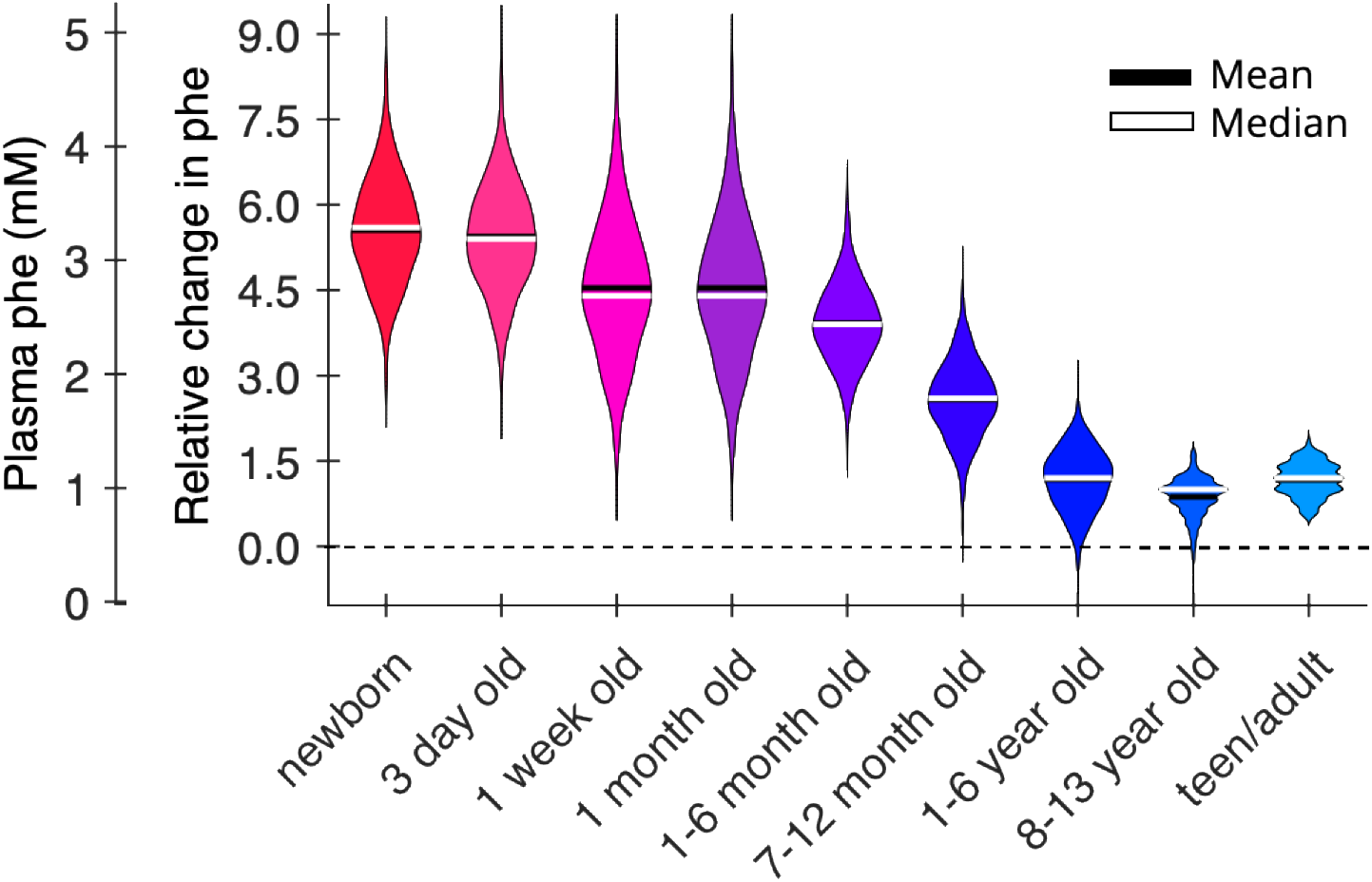
Changes in therapeutic efficacy across different age groups as measured by end-of-day blood phe concentration. Running 1000 simulations of the ALT-CAT model across age groups in which GI physiological properties and meal sizes and feeding schedules vary. The GIT is populated by designating 1% of the bacteria as the living therapeutic and results are represented as a violin plot of end-of-day plasma phe concentration after a 24 h period.

Unfortunately, the physiology of the gut does not lend to metabolic intervention easily, and each age group had plasma phe ubiquitously increase to unsafe levels (Figure 4). As a countering tactical measure, drug formulations for living therapeutics include microencapsulation to survive the stomach (39), surface modification to improve mucoadhesion and increase residence time or cell population (16), bacterial chassis selection to target regional success (40, 41), and enzyme choice and expression to increase specific activity per cell (40, 41). Effectively, the agenda in each case is to increase the level of therapeutic intervention at earlier stages of digestion. This can be simulated in the ALT-CAT model using a coverage multiplier (*αc*) (Equation 13). This is created as the mean of a log-normal distribution to define the living therapeutic population. While it is represented as bacterial concentration in the model, this is a representative parameter of “active therapeutic agent per unit volume of SI” in each compartment, influenced by gastric survival (*G*), availability (*A*), regioselective niches (*R*), surface area (*SA*), transit time (*T_SI_)*, and total microbial population (*M_tot_*) (Figure 5a). By increasing the *α_c_*, the blood plasma phe concentration is noticeably impacted in both children and adults. In the 1- to 6-month-old age group, ALT-CAT model solutions with *αc* = 1, which designates a mean 2.5% of the bacterial population as active, begin to lower plasma phe (Figure 5b). While this is a significant fraction of gut bacteria being replaced with the therapeutic, it corresponds to 10^10^–10^11^ cells, within a reasonable order-of-magnitude for a dosing regimen. As the length of the SI and overall bacterial population is much larger in adults, a lower percent (*α_c_* = 0; mean = 0.8%) is impactful. These comparisons can offer insight into dosing regimen depending on physiology, diet, and bacterial chasses.

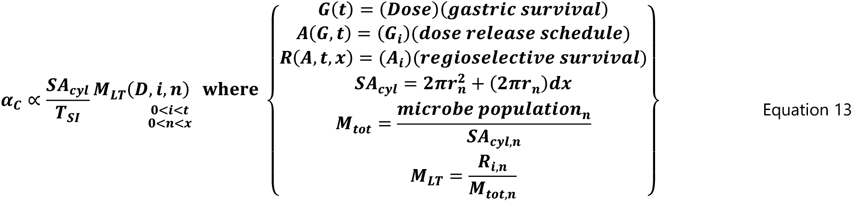

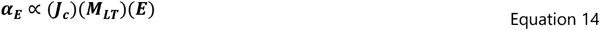

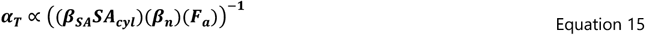

**Figure 5.**
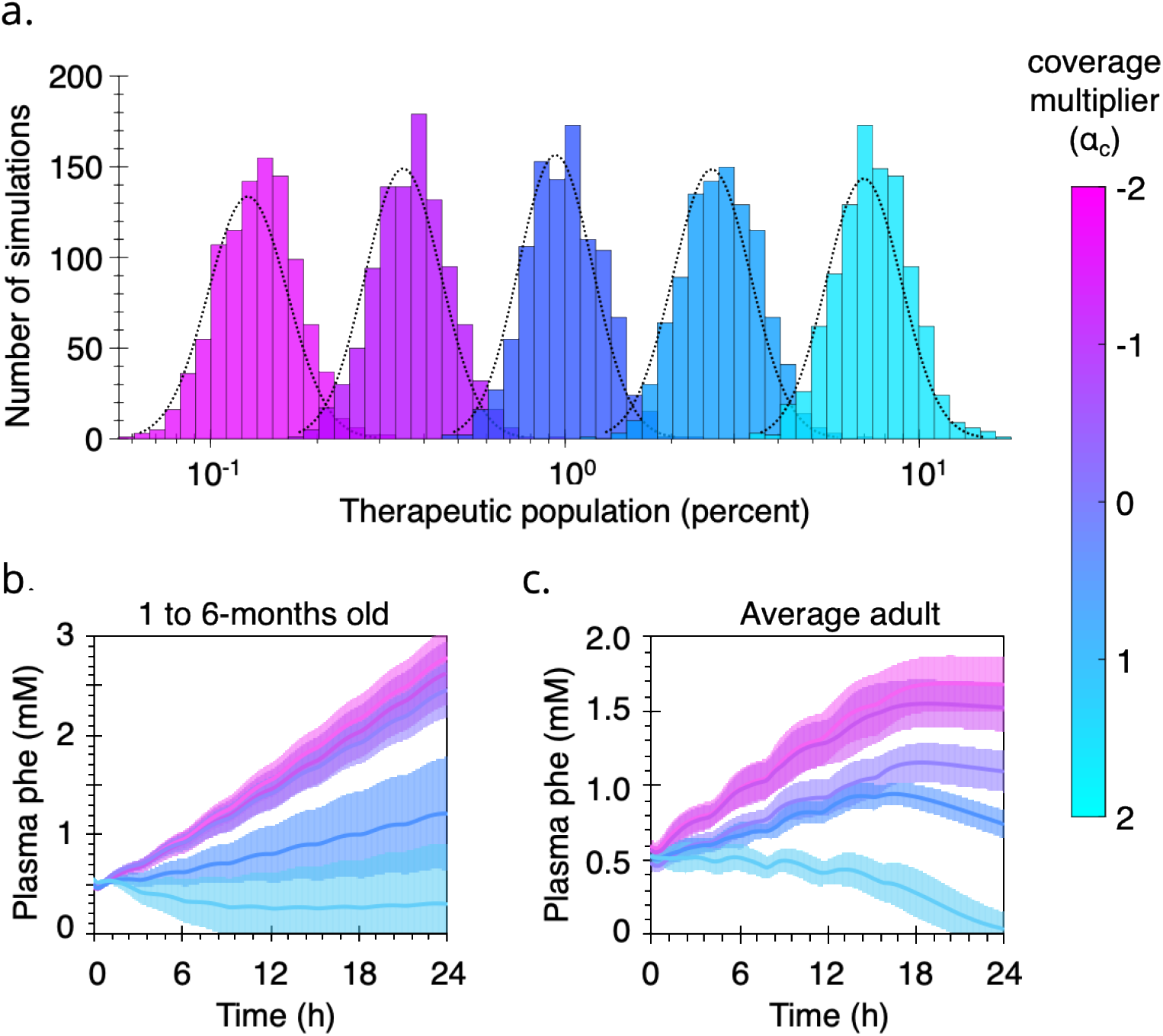
Therapeutic factor impacts on ALT-CAT solutions A PAL-based living therapeutic to treat PKU is highly impacted by the number of bacteria able to degrade phe. (a.) The mean of a lognormal distribution is used to randomly populate the gut within a population range and is represented in the ALT-CAT model as the coverage multiplier (*α_C_*). Varying *α_C_* in the GIT of (b.) 1 to 6-month old babies and (c.) average adults, represented as the blood plasma phe concentration over a 24 h period as mean (line) and standard deviation (shaded region) of 1000 solutions.

As a thought experiment, it is more realistic to control enzyme properties within a bacterial cell than it is to control the many complex environmental interactions bacteria may encounter in the SI. Enzymatic specific activity, whether that be from increased *M_LT_* or higher protein expression (*E*) or turnover (*k_cat_*), can be more informative as a metric in the ALT-CAT model solution space. Thus, similar simulations with each age group were performed using an enzyme multiplier *α_F_*) (Equation 14) applied to PAL rather than microbial population (*α_C_*). Initially, moving from *α_C_* = 0 (~0.8% cells) to *α_C_* = 1 (~2.5% cells) yielded a bacterial population that was efficacious in 1- to 6-month-old babies; this is 3.125-fold change. Reframing this, keeping the bacterial population constant but increasing specific enzyme activity by 3.125-fold (i.e. *α_E_* = 3.125) has an equivalent impact on lowering the end-of-day plasma phe (Figure 6a). That is, enzyme specific activity is proportional to *M_LT_.*

**Figure 6.**
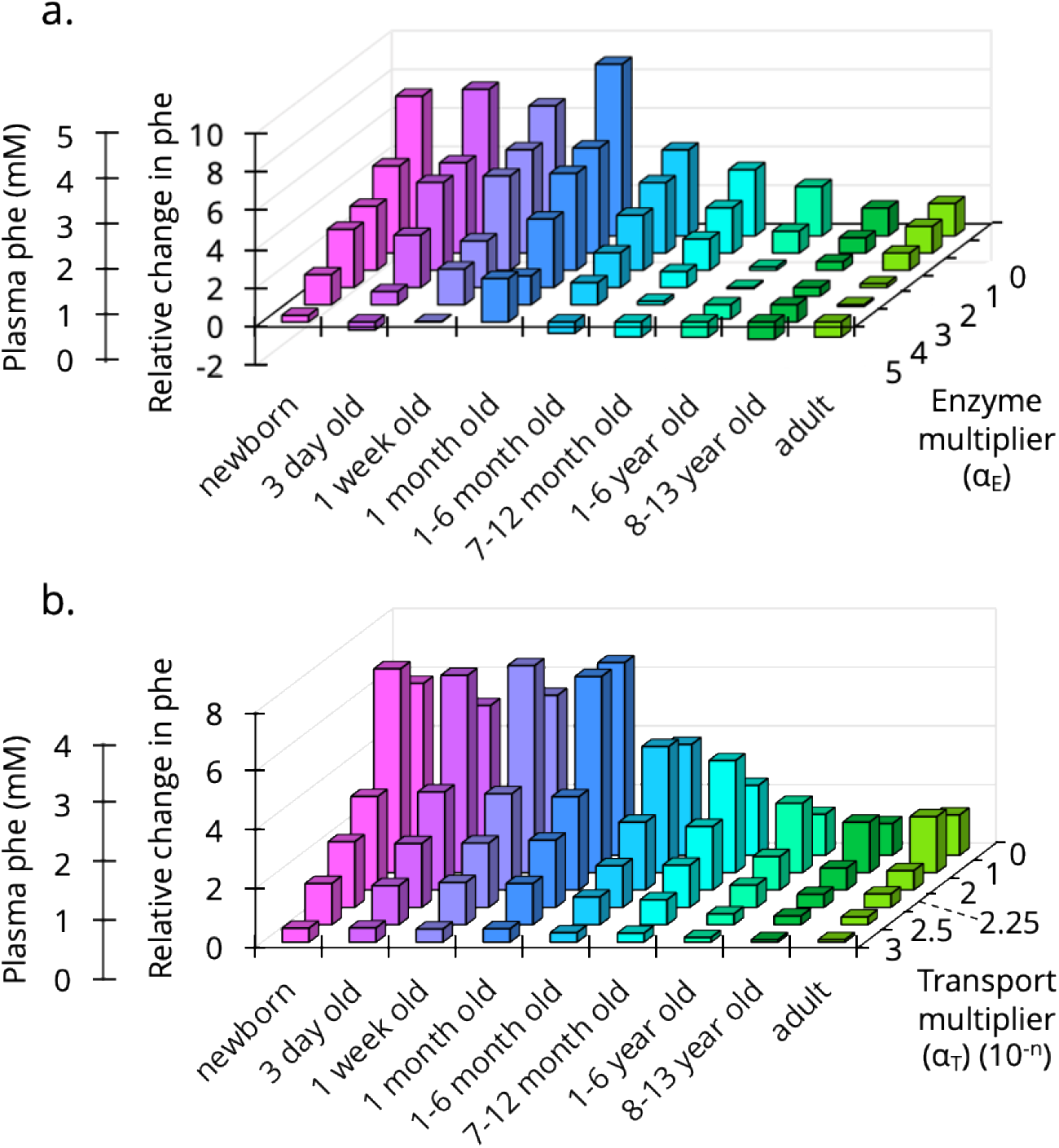
Exploring the solution space of the ALT-CAT model. While keeping the coverage multiplier constant (*α_C_* = 0; ~0.8%), the end-of-day plasma phe concentration was calculated for (a.) increasing PAL specific activity by changing the enzyme multiplier (*α_E_*), and (b.) slowing the absorption of phe by increasing the permeability multiplier (*α_T_*). Each bar is the mean of 100 simulations.

Alternatively, transport of phe across the epithelium is fast, approximately 4.1 × 10^-4^ cm/s (42), and on the high end of typical drugs (42, 43). Slowing *P_eff_* or reducing *β_SA_* or enterorecirculation (*β_n_*) using a transport multiplier (*α_T_*) can offer insight into the weight that transport phenomena impact drug bioavailability (or in the case of PKU, phe absorption) from a living therapeutic (Figure 6b). Lowering the *P_eff_* of phe by 225-fold (*α_T_* = 2.25), improves the treatment 2.3-fold for 1- to 6-month-old babies (Figure 6b). This level of change unlikely to be feasible for the treatment of PKU using LNAA supplementation. However, generalizing beyond phe and treating PKU, this new *P_eff_* ~10^-4^ to 10^-5^ cm/s (i.e. *α_T_* = 2.25) falls within the range of typical small molecule drugs (42, 43). So, considering these trends conceptually as the production of a typical drug with a living therapeutic, if the drug has poor bioavailability (α_T_ = 0), the current-best-reported production (*α_E_* = 1) would require ~2.5% population coverage (*α_c_* = 1). However, improving drug bioavailability by increasing permeability (*α_T_* = 2.25) now only requires ~1% (or 2.5 ÷ 2.3) of the bacterial population to produce the same efficacy, or approximately *α_C_* = 0. Slower absorption requires a lower dose. These model adjustments provide examples as to how tuning different parameters can inform design benchmarks when constructing living therapeutics for different applications.

These three parameters (bacterial population, enzyme activity, and transport) represent avenues of engineering. As such, a combinatorial approach to dial in system- or therapy-specific parameters can be informed by running the ALT-CAT model to set investigative benchmarks and clinical boundaries. Additional drug physiochemical and formulation properties and GI physiological properties can easily be added and assessed using multipliers as explored in Figure 6. While PKU and a PAL-based EST offer a well-suited case study for the development and evaluation of the ALT-CAT model, we present this as a generalized tool for oral drug development and the advancement of living therapeutics as a platform.

## Supplemental Information

The MATLAB files for model can be found at the Nair lab GitHub page: https://github.com/nair-lab/ALT-CAT

## References

1. Amidon GL, Lennernäs H, Shah VP, Crison JR. 1995. A Theoretical Basis for a Biopharmaceutic Drug Classification: The Correlation of in Vitro Drug Product Dissolution and in Vivo Bioavailability. Pharmaceutical Research: An Official Journal of the American Association of Pharmaceutical Scientists 12:413–420.

2. Lipinski CA, Lombardo F, Dominy BW, Feeney PJ. 2012. Experimental and computational approaches to estimate solubility and permeability in drug discovery and development settings. Advanced Drug Delivery Reviews 64:4–17.

3. Turco L, Catone T, Caloni F, Consiglio E di, Testai E, Stammati A. 2011. Caco-2/TC7 cell line characterization for intestinal absorption: How reliable is this in vitro model for the prediction of the oral dose fraction absorbed in human? Toxicology in Vitro 25:13–20.

4. Lennernäs H, Ahrenstedt Ö, Hällgren R, Knutson L, Ryde M, Paalzow LK. 1992. Regional Jejunal Perfusion, a New in Vivo Approach to Study Oral Drug Absorption in Man. Pharmaceutical Research: An Official Journal of the American Association of Pharmaceutical Scientists.

5. Antunes F, Andrade F, Araújo F, Ferreira D, Sarmento B. 2013. Establishment of a triple co-culture in vitro cell models to study intestinal absorption of peptide drugs. European Journal of Pharmaceutics and Biopharmaceutics 83:427–435.

6. Bein A, Shin W, Jalili-firoozinezhad S, Park MH, Kim HJ, Ingber DE. 2018. Microfluidic Organ-on-a-Chip Models of Human Intestine. Cellular and Molecular Gastroenterology and Hepatology 5:659–668.

7. Minekus M, Smeets-Peeters M, Havenaar R, Bernalier A, Fonty G, Marol-Bonnin S, Alric M, Marteau P, Huis In’t Veld JHJ. 1999. A computer-controlled system to simulate conditions of the large intestine with peristaltic mixing, water absorption and absorption of fermentation products. Applied Microbiology and Biotechnology 53:108–114.

8. Basant N, Gupta S, Singh KP. 2016. Predicting human intestinal absorption of diverse chemicals using ensemble learning based QSAR modeling approaches. Computational Biology and Chemistry 61:178–196.

9. Yu LX, Amidon GL. 1999. A compartmental absorption and transit model for estimating oral drug absorption. International Journal of Pharmaceutics 186:119–125.

10. di Meo F, Fabre G, Berka K, Ossman T, Chantemargue B, Paloncýová M, Marquet P, Otyepka M, Trouillas P. 2016. In silico pharmacology: Drug membrane partitioning and crossing. Pharmacological Research 111:471–486.

11. Rowland M, Peck C, Tucker G. 2011. Physiologically-Based Pharmacokinetics in Drug Development and Regulatory Science. Annual Review of Pharmacology and Toxicology 51:45–73.

12. Willmann S, Thelen K, Lippert J. 2012. Integration of dissolution into physiologically-based pharmacokinetic models III: PK-Sim. Journal of Pharmacy and Pharmacology 64:997–1007.

13. Marsousi N, Desmeules JA, Rudaz S, Daali Y. 2018. Prediction of drug-drug interactions using physiologically-based pharmacokinetic models of CYP450 modulators included in Simcyp software. Biopharmaceutics and Drug Disposition 39:3–17.

14. Gobeau N, Stringer R, de Buck S, Tuntland T, Faller B. 2016. Evaluation of the GastroPlus™ Advanced Compartmental and Transit (ACAT) Model in Early Discovery. Pharmaceutical Research 33:2126–2139.

15. Agoram B, Woltosz WS, Bolger MB. 2001. Predicting the impact of physiological and biochemical processes on oral drug bioavailability. Advanced Drug Delivery Reviews 50:S41–S67.

16. Mays ZJ, Nair NU. 2018. Synthetic biology in probiotic lactic acid bacteria: At the frontier of living therapeutics. Curr Op Biotechnol 53:224–231.

17. Bober JR, Beisel CL, Nair NU. 2018. Synthetic Biology Approaches to Engineer Probiotics and Members of the Human Microbiota for Biomedical Applications 277–300.

18. Waller MC, Bober JR, Nair NU, Beisel CL. 2017. Toward a genetic tool development pipeline for host-associated bacteria. Curr Opin Microbiol 38:156–164.

19. Mimee M, Citorik RJ, Lu TK. 2016. Microbiome therapeutics — Advances and challenges. Advanced Drug Delivery Reviews 105:44–54.

20. Isabella VM, Ha BN, Castillo MJ, Lubkowicz DJ, Rowe SE, Millet YA, Anderson CL, Li N, Fisher AB, West KA, Reeder PJ, Momin MM, Bergeron CG, Guilmain SE, Miller PF, Kurtz CB, Falb D. 2018. Development of a synthetic live bacterial therapeutic for the human metabolic disease phenylketonuria. Nat Biotechnol 36:857–867.

21. Chang S, Ming T, Bourget L, Lister C. 1995. A new theory of enterorecirculation of amino acids and its use for depleting unwanted amino acids using oral enzyme-artificial cells, as in removing phenylalanine in phenylketonuria. Artificial Cells, Blood Substitutes, and Biotechnology 23:1–21.

22. West KA, Perreault M, Kurtz CB, Wagner DA, Charbonneau MR, Dagon Y, Degar AJ, Kotula JW, Millet YA, Brennan AM, Miller PF, Puurunen MK, Denney WS, Isabella VM, Antipov E. 2019. An engineered E. coli Nissle improves hyperammonemia and survival in mice and shows dose-dependent exposure in healthy humans. Sci Transl Med 11:eaau7975.

23. Rhimi M, Bermudez-Humaran LG, Huang Y, Boudebbouze S, Gaci N, Garnier A, Gratadoux JJ, Mkaouar H, Langella P, Maguin E. 2015. The secreted l-arabinose isomerase displays anti-hyperglycemic effects in mice. Microbial Cell Factories 14:204.

24. Lin Y, Krogh-Andersen K, Hammarström L, Marcotte H. 2017. Lactobacillus delivery of bioactive interleukin-22. Microbial Cell Factories 16:148.

25. Zhang B, Li A, Zuo F, Yu R, Zeng Z, Ma H, Chen S. 2016. Recombinant Lactococcus lactis NZ9000 secretes a bioactive kisspeptin that inhibits proliferation and migration of human colon carcinoma HT-29 cells. Microbial Cell Factories 15:102.

26. Yu LX, Crison JR, Amidon GL. 1996. Compartmental transit and dispersion model analysis of small intestinal transit flow in humans. Int J Pharm 140:111–118.

27. Yokrattanasak J, de Gaetano A, Panunzi S, Satiracoo P, Lawton WM, Lenbury Y. 2016. A simple, realistic stochastic model of gastric emptying. PLoS ONE 11:1–15.

28. Locatelli I, Mrhar A, Bogataj M. 2009. Gastric emptying of pellets under fasting conditions: A mathematical model. Pharmaceutical Research 26:1607–1617.

29. Macheras P, Karalis V, Valsami G. 2013. Keeping a critical eye on the science and the regulation of oral drug absorption: A review. Journal of Pharmaceutical Sciences 102:3018–3036.

30. Hu M, Borchardt RT. 1992. Transport of a large neutral amino acid in a human intestinal epithelial cell line (Caco-2): uptake and efflux of phenylalanine. BBA - Molecular Cell Research 1135:233–244.

31. Hidalgo IJ, Borchardt RT. 1990. Transport of a large neutral amino acid (phenylalanine) in a human intestinal epithelial cell line: Caco-2. BBA - Biomembranes 1028:25–30.

32. Walter J, Ley R. 2011. The Human Gut Microbiome: Ecology and Recent Evolutionary Changes. Annu Rev Microbiol 65:411–429.

33. Douillard FP, de Vos WM. 2014. Functional genomics of lactic acid bacteria: From food to health. Microb Cell Fact 13:S8.

34. Helander HF, Fändriks L. 2014. Surface area of the digestive tract-revisited. Scandinavian Journal of Gastroenterology 49:681–689.

35. Zhang Y, Jia X, Wang L, Liu J, Ma G. 2011. Preparation of Ca-alginate microparticles and its application for phenylketonuria oral therapy. Industrial and Engineering Chemistry Research 50:4106–4112.

36. Matalon R, Michals-Matalon K, Bhatia G, Burlina AB, Burlina AP, Braga C, Fiori L, Giovannini M, Grechanina E, Novikov P, Grady J, Tyring SK, Guttler F. 2007. Double blind placebo control trial of large neutral amino acids in treatment of PKU: Effect on blood phenylalanine. Journal of Inherited Metabolic Disease 30:153–158.

37. Mariotta L, Ramadan T, Singer D, Guetg A, Herzog B, Stoeger C, Palacín M, Lahoutte T, Camargo SMR, Verrey F. 2012. T-type amino acid transporter TAT1 (Slc16a10) is essential for extracellular aromatic amino acid homeostasis control. Journal of Physiology 590:6413–6424.

38. Kim DK, Kanai Y, Chairoungdua A, Matsuo H, Cha SH, Endou H. 2001. Expression Cloning of a Na + -independent Aromatic Amino Acid Transporter with Structural Similarity to H+/Monocarboxylate Transporters. Journal of Biological Chemistry 276:17221–17228.

39. Mays ZJS, Chappell TC, Nair NU. 2020. Quantifying and Engineering Mucus Adhesion of Probiotics. ACS Synthetic Biology acssynbio.9b00356.

40. Jendresen CB, Stahlhut SG, Li M, Gaspar P, Siedler S, Förster J, Maury J, Borodina I, Nielsen AT. 2015. Highly active and specific tyrosine ammonia-lyases from diverse origins enable enhanced production of aromatic compounds in bacteria and Saccharomyces cerevisiae. Applied and Environmental Microbiology 81:4458–4476.

41. Wu B, Szymanski W, Heberling MM, Feringa BL, Janssen DB. 2011. Aminomutases: Mechanistic diversity, biotechnological applications and future perspectives. Trends in Biotechnology 29:352–362.

42. Dahlgren D, Roos C, Sjögren E, Lennernäs H. 2015. Direct in Vivo Human Intestinal Permeability (Peff) Determined with Different Clinical Perfusion and Intubation Methods. Journal of Pharmaceutical Sciences 104:2702–2726.

43. Wolk O, Markovic M, Porat D, Fine-Shamir N, Zur M, Beig A, Dahan A. 2019. Segmental-Dependent Intestinal Drug Permeability: Development and Model Validation of In Silico Predictions Guided by In Vivo Permeability Values. Journal of Pharmaceutical Sciences 108:316–325.

